## Supplementary figures and images for "The Drosophila cancer-germline, head-to-head gene pair *TrxT* and *dhd* is dispensable for normal brain development but plays a major role in *l(3)malignant brain tumour* growth"

### Figure S1

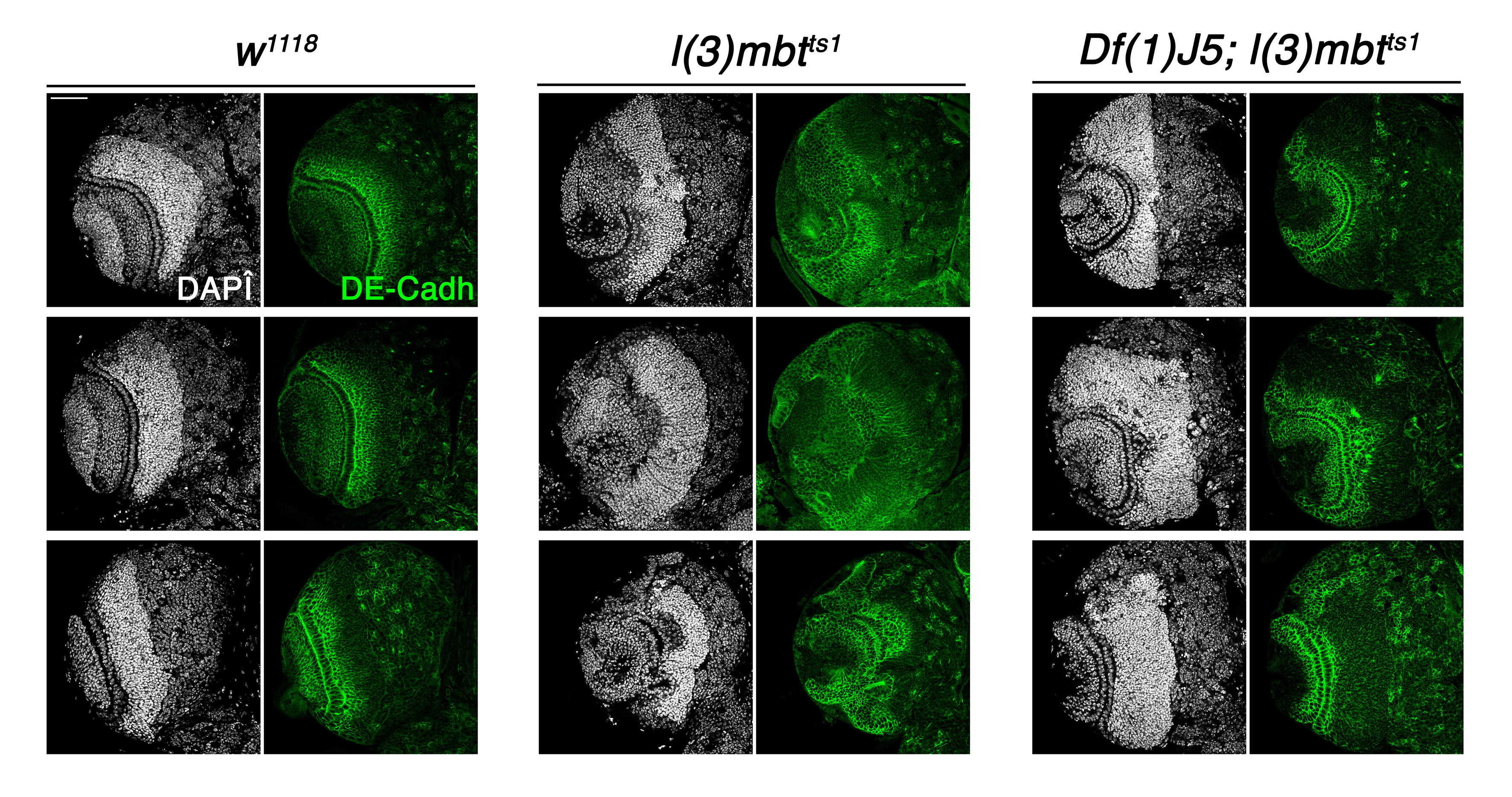

### Figure S2

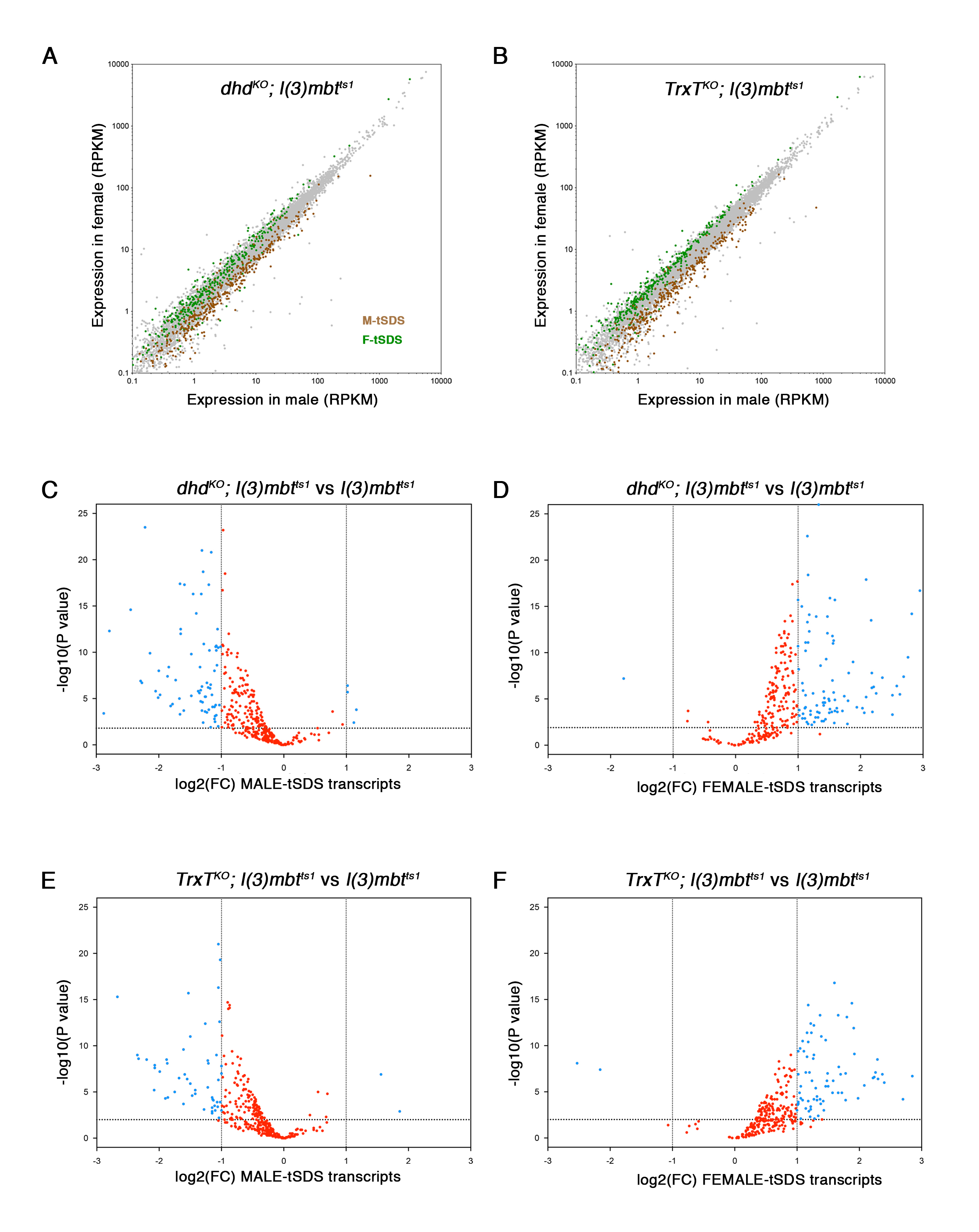
